# Early Stages of Misfolding of PAP248-286 at two different pH values: An Insight from Molecular Dynamics Simulations

**DOI:** 10.1101/2022.06.01.494297

**Authors:** Nikhil Agrawal, Emilio Parisini

**Affiliations:** Latvian Institute of Organic Synthesis, Aizkraukles 21, LV, 1006, Riga, Latvia; College of Health Sciences, University of KwaZulu-Natal, Durban, South Africa

**Keywords:** HIV, PAP248-286, MD Simulations

## Abstract

PAP248-286 peptides, which are highly abundant in human semen, aggregate and form amyloid fibrils that enhance HIV infection. Previous experimental studies have shown that the infection-promoting activity of PAP248-286 begins to increase well before amyloid formation takes place and that pH plays a key role in the enhancement of PAP248-286-related infection. Hence, understanding the early stages of misfolding of the PAP2482-86 peptide is crucial. To this end, we have performed 60 independent MD simulations for a total of 24 μs at two different pH values (4.2 and 7.2). Our data shows that misfolding of the PAP248-286 peptide is a multistage process and that the first step of the process is a transition from an “I-shaped” structure to a “U-shaped” structure. We further observed that the structure of PAP248-286 at the two different pH values shows significantly different features. At pH 4.2, the peptide has less intra-molecular H-bonds and a reduced α-helical content than at pH 7.2. Moreover, differences in intra-peptide residues contacts are also observed at the two pH values. Finally, free energy landscape analysis shows that there are more local minima in the energy surface of the peptide at pH 7.2 than at pH 4.2. Overall, the present study elucidates the early stages of misfolding of the PAP248-286 peptide at the atomic level, thus possibly opening new avenues in structure-based drug discovery against HIV infection.

## Introduction

HIV has infected more than 75 million people since its inception and, so far, has claimed the lives of more than 36 million people worldwide^1,2^. Human semen is one of the most important vectors for HIV transmission and women are at high risk of HIV through unprotected vaginal intercourse^3,4–5^. The one very crucial natural factor that has been identified to play a key role in HIV transmission is fragments of the prostatic acidic phosphatase (PAP) protein^6^. The region of PAP from amino acid number 248 to 286 (PAP248-286) is naturally present in high quantity in the human semen. Studies have reported that the PAP248-286 peptide could be responsible for an increase in the rate of infection of HIV by up to 400000-fold^7,8^. PAP248-286 peptides aggregate and form amyloid fibrils termed Semen-derived enhancers of viral infection (SEVI). The highly cationic nature of the SEVI fibrils facilitates their binding to the anionic membranes of HIV virions and cells, which leads to an increase in virion cell attachment and fusion^9,10,11^. In the recent times, several different approaches have been taken to reduce the PAP248-286-mediated rate of HIV infection. These include (i) inhibition of the transformation of PAP248-286 monomers into amyloids, iii) hindering of the cleavage of the PAP248-286 peptide from the PAP protein, iii) perturbation of the structure of the SEVI fibrils, and iv) neutralization of the overall charge of the SEVI fibrils. Out of the above-mentioned approaches, inhibition of early stages of misfolding of PAP248-286 monomers could be very promising, as it has been reported to similar amyloidogenic peptides that are related to Alzheimer’s and Parkinson’s diseases^11–12^. While the biological activity of PAP248-286 is mostly reported to its aggregated amyloid form, studies have shown that the monomeric form of PAP248-286 is also biologically active^6,13^.

Previous studies using in vitro and in vivo experiments reported that PAP248-286 shows HIV viral enhancement activity mainly at the acidic vaginal environment (pH ~4.2)^7,14^. Brender^14^ *et. al.* suggested that the high activity of PAP248-286 in the acidic vaginal environment may be due to its two histidine residues, which are likely to be charged at the acidic pH ~4.2, but not at the neutral pH ~7.2. Interestingly, other than showing HIV infection enhancement at acidic pH, the PAP248-286 peptide has also been reported to display antimicrobial activity through bacterial agglutination at neutral pH^15,13^.

Several experimental studies, mostly using the NMR technique, have been performed to understand the monomeric structure of the full-length PAP248-286 peptide or short regions thereof in different environments. It has been reported that in aqueous and SDS micelles environments the structure of the PAP248-286 peptide is highly disordered and can be divided into three parts: i) a N-terminal disordered region, ii) a stable central part, which forms an α-helix, iii) a C-terminal flexible region, which adopts a 3_10_/α-helix conformation^16,17,18,19,20,21^. In trifluoroethanol (TFE), PAP248-286 features a largely α-helical conformation. Indeed, in 30% TEF, almost all the central part of the peptide adopts a helical conformation while in 50% TEF the whole peptide, with the exception of its C-terminal end, displays a α-helical conformation. The conformation of PAP248-286 in 50% TEF solution is similar to the PAP248-286 conformation in the crystal structure of the full-length PAP protein (PDB id : 1CVI)^16^.

Misfolding and aggregation of proteins to form amyloids is a well-known phenomenon, particularly but not exclusively in the context of neurodegenerative diseases such as Alzheimer’s disease (AD)^22,23,24^ and Parkinson’s disease (PD)^25^. In fact, there are almost 50 other human diseases that are related to the formation of amyloid^26^. Understanding the early stages of misfolding of amyloid-forming proteins may help identify amino acid residues that play a key role in the process, and aid the design of potential aggregation inhibitors^27^. Molecular dynamics (MD) simulations are widely used to investigate the misfolding and aggregation process of amyloid proteins. MD simulations provide an atomic-level insight into the transformation of α-helix-rich structures into a disordered or β-rich structures that is currently not attainable by experimental techniques^28^. In recent times, several MD simulations studies have been conducted on the PAP248-286 peptide; however, all these studies were done to investigate the binding of small molecules to PAP248-286^29,30^ or its interaction with biological membrane lipids^31,13^. To the best of our knowledge, no MD simulation study has been conducted so far to investigate the early stages of misfolding of PAP248-286 at the acidic pH characteristic of the vaginal environment and at neutral pH. The overall aim of the present study is to investigate and compare the mechanism of the early stages of misfolding of PAP248-286 at two different pH values. To this end, we have performed 60 independent all-atoms MD simulations of PAP248-286 at pH 4.2 and at pH 7.2 in explicit water environment.

## Methods

In the present study, we used the NMR structure of PAP248-286 (PDB id: 2L77)^16–17^ for our molecular dynamics simulations. This PAP248-286 peptide structure closely resembles the conformation of the same amino acid portion in the crystal structure of the full-length PAP (PDB id: 1CVI)^32^. Residue numbering of PDB id: 2L77 starts from 1 and ends at 39. At pH 4.2, His3 and His23 of PAP248-286 are protonated, leading to an overall charge of +8 on the peptide, compared to a +6 charge at pH 7.2 (Figure 1A, B). We used the H^++^ server^33^ to add hydrogen atoms to PAP248-286: at pH 4.2 to account for the acidic vaginal environment and at pH 7.2 to represent the physiological environment. In the present study, we used the charmm36m force field^34^, which is an updated version of the charmm36 force field^35^. The charmm36m force field has shown improved accuracy in generating conformational ensembles for intrinsically disordered peptides and proteins^34,36^. Our simulation system contains a total of 12031 water molecules at pH 4.2 and 12039 water molecules at pH 7.2. To neutralize the overall charge, 8 Cl^-^ ions were added in the pH 4.2 system and 6 Cl^-^ ions were added in the pH 7.2 system. Both systems were energy minimized using the steepest descent algorithm, followed by two sequential 500 picoseconds (ps) equilibration simulations using the canonical (NVT) and isobaric-isothermic (NPT) ensemble, and production simulations were performed using the NPT ensemble. During energy minimization and the first equilibration step, restrains of 400 and 40 kJ mol^-1^ nm^-2^ were employed on backbone and sidechain atoms of PAP248-286, respectively. In the second equilibration step, restrains were maintained only on backbone atoms. In the production runs, no restrains were employed on protein atoms. The LINCS algorithm was used for constraining the bond lengths from heavy atoms to hydrogen and the SETTLE algorithm was employed for water molecule bond length constraints. The Particle mesh Ewald (PME) algorithm was used for long-range electrostatic interactions, while for short-range van der Waals (vdW) and columbic interactions a cut-off of 12 Å was used. The Nose-Hoover algorithm was used for temperature coupling and the Parrinello-Rahman algorithm was employed for pressure coupling. A 1 ps time constant was used for both temperature and pressure coupling. All production simulations were performed at a pressure of 1 bar and at a temperature of 310.15 K. A leap-frog algorithm was used for integrating Newton’s equations of motion. A total of 60 simulations were performed (30 simulations at each pH value) using different initial velocities randomly generated by the GROMACS package.

**Figure 1.**
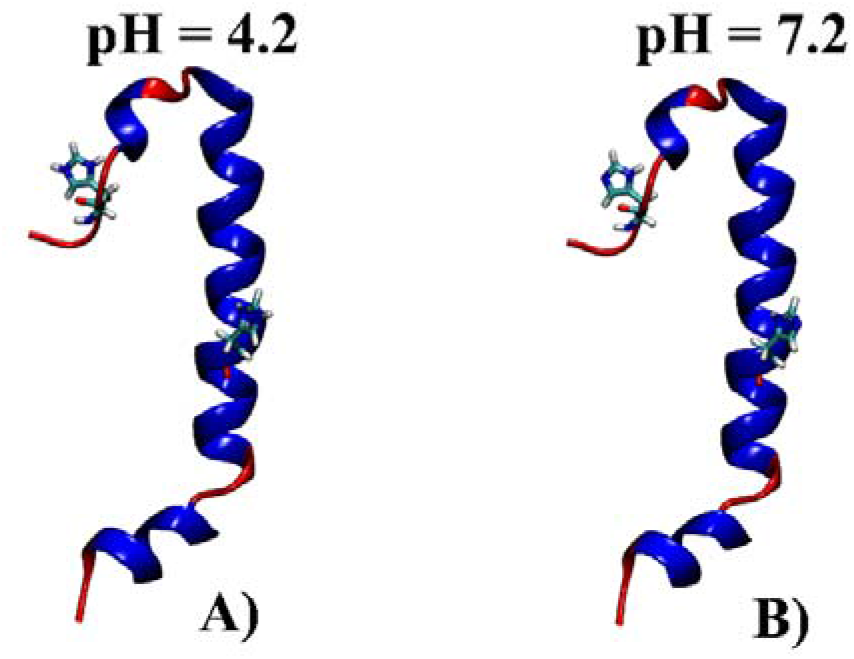
A) PAP248-286 structure with both histidine residues protonated (pH 4.2), b) PAP248-286 NMR structure (PDB id = 2L77) at pH 7.2, with both histidine residues deprotonated. The helical portion of the peptide is shown in blue color, while the unstructured portions of the peptide are shown in red color.

### Analysis Details

Time evolution of backbone RMSD of PAP248-286 was calculated with respect to the NMR structure. Hydrogen (H) bonds analysis was performed using the gromacs^37^ hbond program and employing the default parameters (donor and acceptor atoms distance cut-off 3.5 Å, and hydrogen-donor-acceptor angle cut-off 30°). The transition of PAP248-286 from an “I-shaped” structure to a “U-Shaped” structure was calculated using the gromacs^37^ angle program and the Cα atoms of Val15, Thr28 and Leu36 were used for angle calculation. Residue contact maps, total interaction time, interaction types, and native contacts were calculated with the CONtact ANalysis (CONAN) tool^38^ using backbone atoms and rcut was set at 10 Å, rinter and rhigh-inter were set at 5 Å as used in the reference paper of CONAN tool^38^. Above the rcut value, any pair of residues that does not have at least one atom in this range is not considered. Contact is formed between a pair of residues under rinter and interactions break above rhigh-inter cutoff value^39^. The interaction type lifetime cut-off was set to 0.50 (representing only interactions that are physically relevant and last at least 50% of the trajectory, i.e. 6 μs or above). Free energy landscape (FEL) was constructed with the gromacs^37^ sham program usi ng RMSD (backbone atoms) and the fraction of native contacts (Q) as two order parameters. The initial NMR structure of PAP248-286 was used as reference for RMSD and Q calculation. BitClust program^40^ was used for the clustering of PAP248-286 structures with a RMSD cut-off of 5Å. Secondary structure analysis of PAP248-286 was performed using the Dictionary of Secondary Structure of Proteins program (DSSP)^41^. For analysis of RMSD boxplot, ROG boxplot H-Bonds length distribution, H-Bonds angle distribution, residue contact maps, total interaction time, interaction types, native contacts, FEL and cluster analyses all 30 trajectories were combined to make a single trajectory (12 μs) at both pH values.

## Results and discussion

### Effect of pH on conformation transition of PAP248-286

RMSD is the most used method to provide a quantitative measure of the similarity between two superimposed atomic coordinates^42^. To investigate the effect of pH on the change in conformation of the PAP248-286 peptide we have plotted the time evolution of its backbone RMSD for all trajectories with respect to its initial structure, whereas to visualize the distribution of RMSD values we have used boxplots (Figure. 2). Time evolution of backbone RMSD shows that large conformational changes take place in PAP248-286 at both pH values. However, at pH 4.2 we observe a lager and a faster conformational change in the structure of the peptide compared to pH 7.2. (Figure 2A, B).

**Figure 2.**
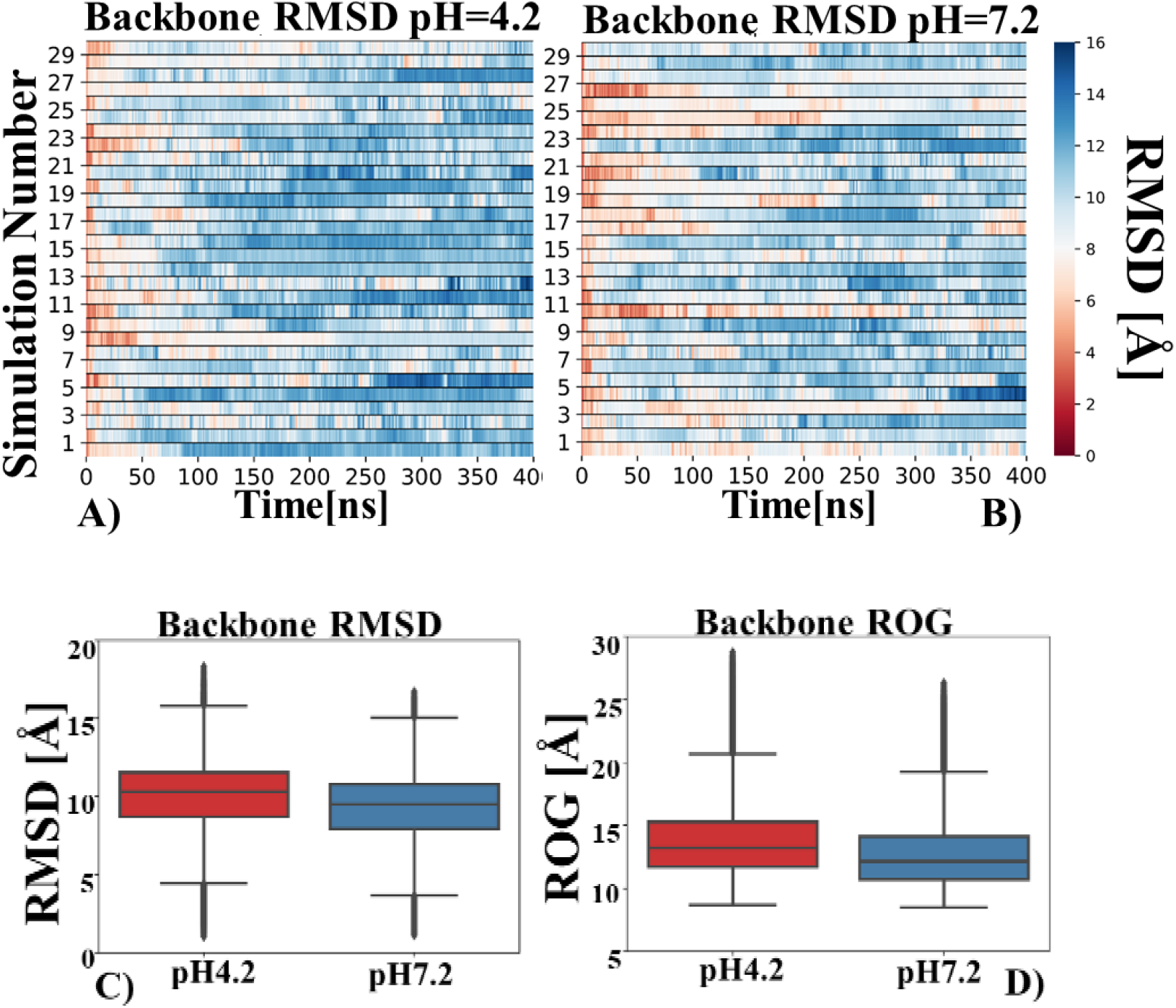
A) Time evolution of the backbone RMSD of PAP248-286 at pH 4.2. B) Time evolution of the backbone RMSD of PAP248-286 at pH 7.2. More blue color represents high RMSD value. C) Backbone RMSD boxplots at pH 4.2 and pH 7.2. D) Radius of gyration boxplot at pH 4.2 and pH 7.2.

Figure 2C provides a further insight into the RMSD distribution, showing that at pH 4.2 the majority of the RMSD values are within the range ~8.8 Å to ~11.8 Å while at pH 7.2 they are mostly comprised between ~7.8 Å and ~10.8 Å. Finally, to assess PAP248-286 change in compactness, we calculated the radius of gyration (ROG) of the peptide (Figure 2D). ROG values at pH 4.2 are mainly distributed between ~11 Å to ~14.8 Å, while at pH 7.2 they are between ~10 Å to ~13.8 Å. Overall, this quantitative data suggests that, relative to its initial conformation, there are more conformational changes in the PAP248-286 peptide at pH 4.2 than at pH 7.2.

### Intra-peptide hydrogen bonding

Hydrogen bonds play a key role in protein/peptide stability as they provide the majority of directional interactions and help protein/peptide to adopt stable secondary structures such as α-helices and β-sheets^43^. Loss of hydrogen bonds often leads to enhanced flexibility in protein/peptide and may promote transition to unfolded/misfolded state^44^. To evaluate the effect of pH on the change of intra-molecular hydrogen bonds in PAP248-286, we have calculated time evolution–H-bonds at the two pH values (Figure 3). Interestingly, while we see a loss of intra-molecular hydrogen bonds at both pH values, we observe that more H-bonds are lost at pH 4.2 and that this occurs in more trajectories (Figure 3A). To quantify the total number of hydrogen bonds, we have calculated an average number of hydrogen bonds per timeframe for the last 100 ns for each trajectory and then we have calculated its average value and standard deviation over the whole set of runs. At pH 4.2, there are ~14.01 ± 3.47 hydrogen bonds per frame in the last 100 ns, while at pH 7.2 their number changes to ~16.32 ± 3.32. Hence, on average in the last 100 ns there are ~2.31 more hydrogen bonds per timeframe of trajectory at pH 7.2 than at pH 4.2. To observe the effect of pH on the lengths and the bond angles of the hydrogen bonds, we have plotted the distribution of these two parameters for all the trajectories combined at both pH values. Figure 4 shows that there are noticeable differences in the distribution of hydrogen bond lengths and relative frequency of bond angle. At pH 7.2, distribution frequency of bond length at around bond length value ~2.95 to 3.0 Å is higher than at pH 4.2. Additionally, relative frequencies of hydrogen bond angles > 10° is also higher at pH 7.2. Taken together, this data suggests that at pH 4.2 the peptide is more flexible and unstable than at pH 7.2.

**Figure 3.**
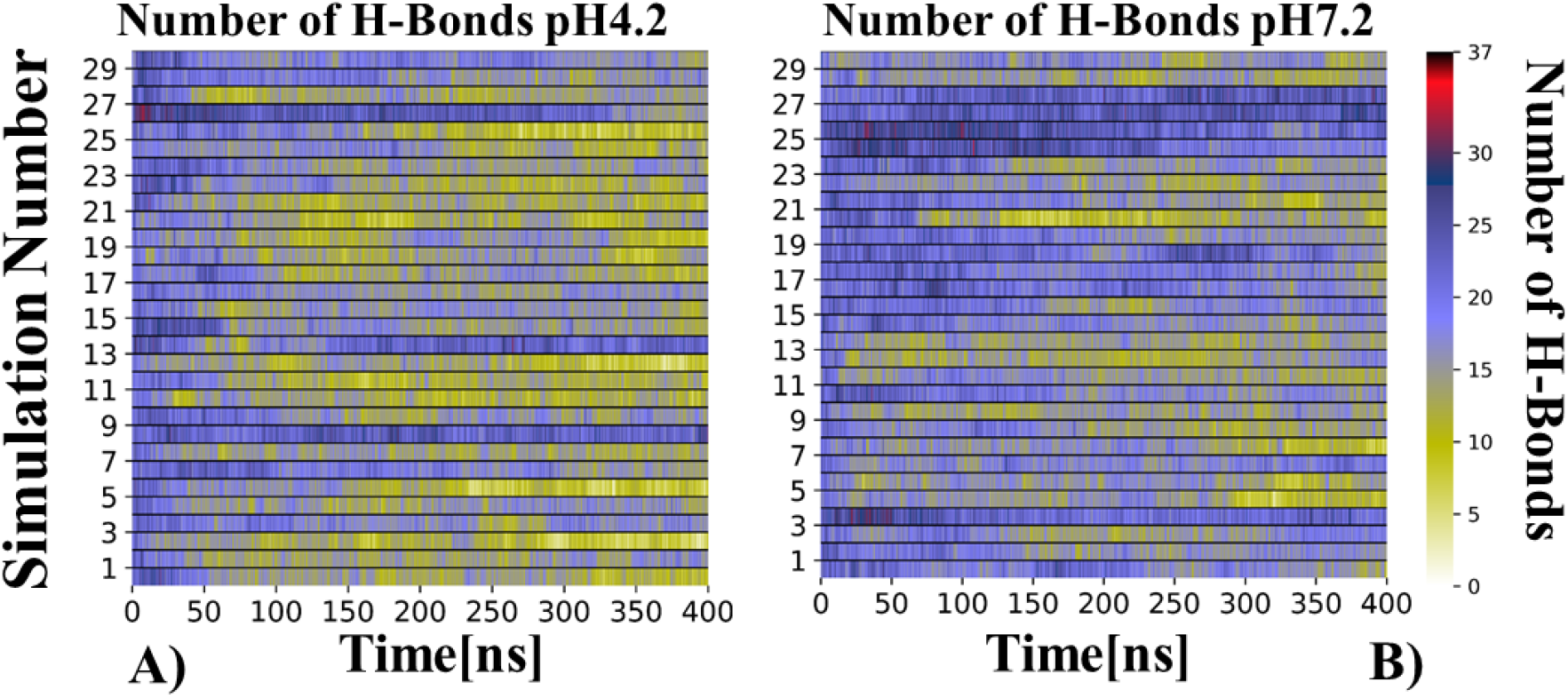
A) Time evolution of the number of hydrogen bonds at pH 4.2. B) Time evolution of the number of hydrogen bonds at pH 7.2. Blue color indicates a higher number of hydrogen bonds while yellow color indicates a lower number of hydrogen bonds.

**Figure 4.**
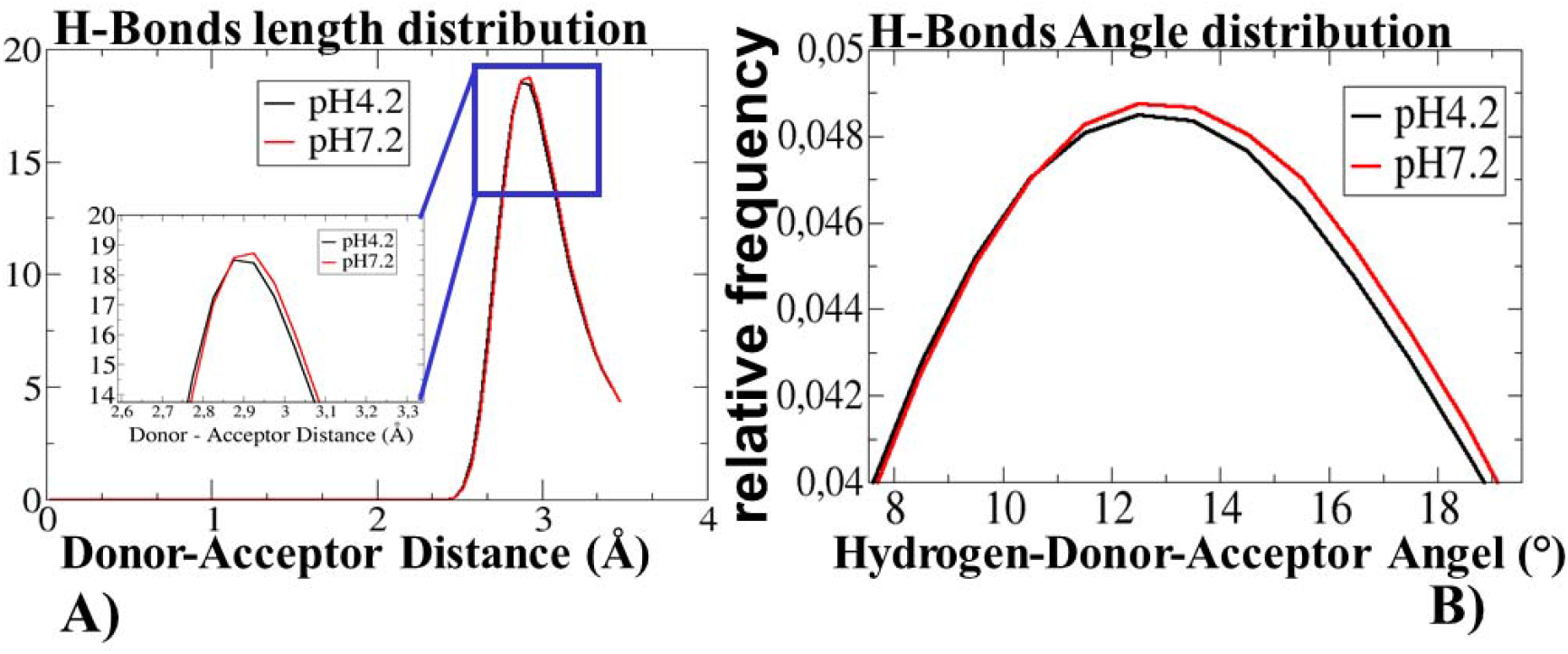
A) Distribution of hydrogen bond length at pH 4.2 and pH 7.2, B) Relative frequency of hydrogen bond angles at pH 4.2 and pH 7.2.

### Transition of PAP248-286 towards a “U-shaped” conformation at pH 4.2 and pH 7.2

The literature has reported that the first step in the multistage process of amyloid fibril formation is the misfolding of the monomeric peptide and that misfolded monomers can induce conformational changes in other monomeric peptides^45,46^. Previous MD simulations studies have suggested that misfolding in an amyloid monomer starts with the formation of a turn, such as for instance in the α-Synuclein (αS) peptide, where turn formation takes place in the four-residue region Thr44-Gly47^47^, and in the Aβ peptide, where the most prevalent region for turn formation is between Val24 and Lys28^27^. In amyloid monomer peptides, turn formation causes a change in conformation from an “I-shaped” structure to a “U-shaped” structure. To investigate the transition of the PAP248-286 peptide structure from “I-Shaped” to “U-shaped” (Figure 5) and to understand the effect of pH on this transition, we have calculated an angle of bending (see the method section for more details) for all simulation trajectories. Bending angle ≤ 90° corresponds to a “U-shaped” topology. We observed that, at pH 4.2, turn formation takes place in all 30 trajectories and that in 27 of them (bending angle ≤ 90°’ green color in Figure 5A) it remains until the end of the simulation time. At pH 7.2, turn formation also takes place in all 30 trajectories, remaining until the end of the simulation time in 23 of them (Figure 5B). The model of the PAP248-286 structure at different time points from one of the representative simulations at both pH values is shown in Figure 6.

**Figure 5.**
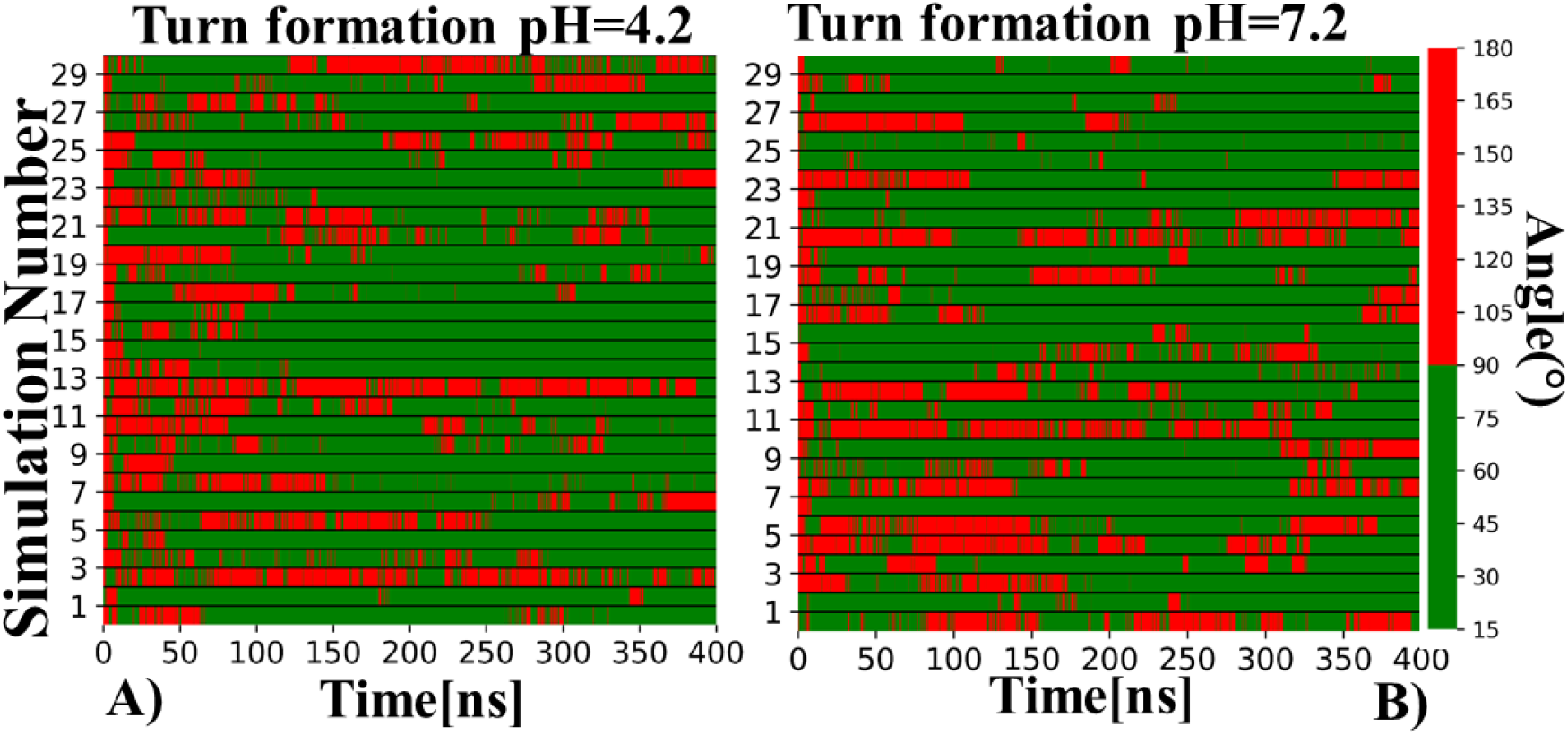
Time evolution of turn formation in the PAP248-286 peptide at A) pH 4.2 and B) pH 7.2.

**Figure 6.**
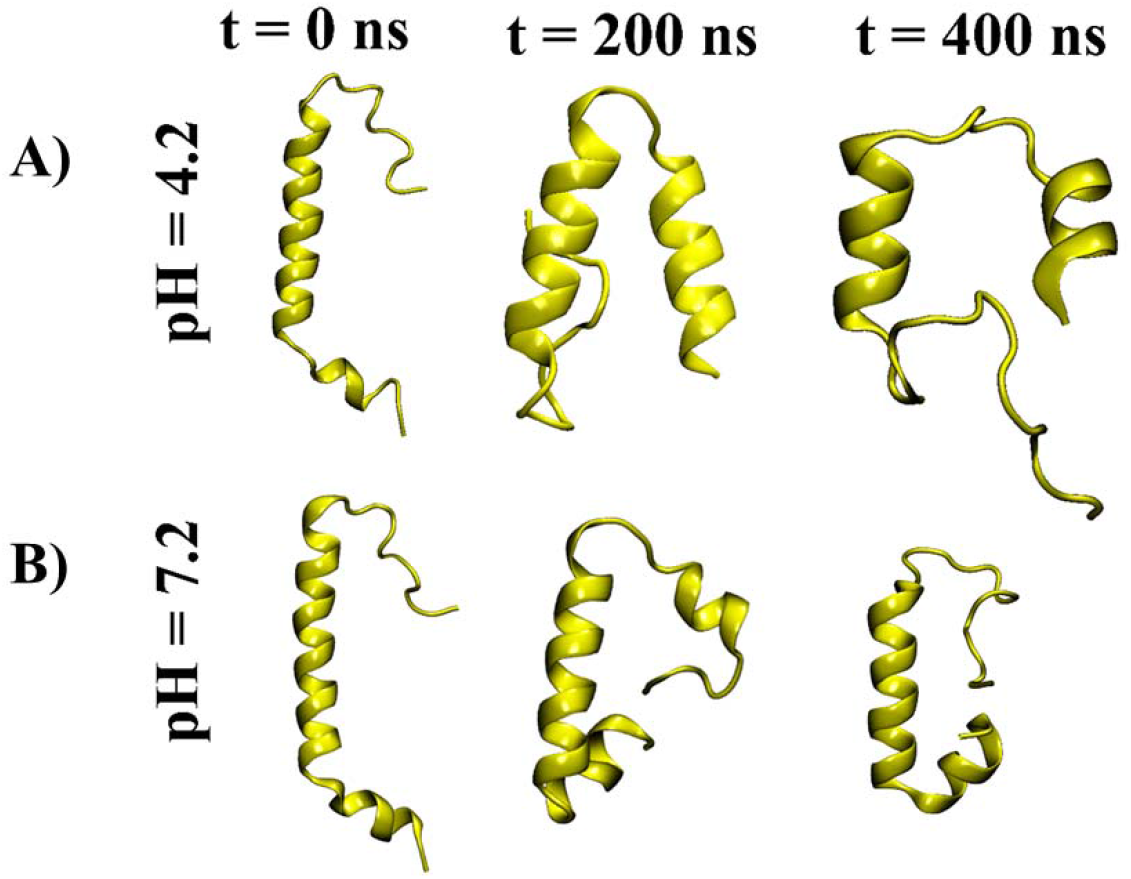
Model structures of PAP248-286 from one of the representative simulations at three different time points A) at pH = 4.2 and B) at pH = 7.2.

To analyze the region involved in turn formation, we performed a visual inspection of all trajectories at the two different pH values. At both pH values, turn formation is most prevalent in the residue region 26RATQ29, albeit with a significantly different frequency. At pH 4.2 and at pH 7.2, turn formation takes place in this region in 21 and in 15 runs, respectively. Turn formation is also observed in the region 29QIP31 at both pH values. Other regions are specific to pH 4.2 (24MK25, 24MKR26, 23HMKR26, 30IPS32, and 23DMK25) or to pH 7.2 (28TQ29, 27ATQ29, 30IPS32, 30IP31, 24MKRA27, 25KRA27, 28TQI30). Overall, this data suggests that there is more variability in turn formation region at pH 7.2 than at pH 4.2. C-terminal residues between 24 and 32 (MKRATQIPS) are likely to be involved in turn formation at both pH values (supplementary Table 1).

### Changes in secondary structure in PAP248-286

Previous studies have shown that the secondary structure of a protein/peptide is affected by external factors such as temperature, pressure and pH^48,49^. To see the effect of pH on the secondary structure of the PAP248-286 peptide, we have calculated the time evolution of the secondary structure of each trajectory at pH 4.2 (Figure S1) and at pH 7.2 (Figure S2). Time evolution of secondary structure data reveals that, at pH 4.2, PAP248-286 loses more of its helical structure than at pH 7.2 (Figure S2). To see a distribution of α-helix and coil content in all trajectories we have plotted boxplots of the percentage of α-helix and coil content for the last 100 ns (300-400 ns) at both pH values. (Figure 7). The plot shows that that α-helix content in PAP248-286 is mostly in the range between ~22% and ~28% at pH 4.2 (Figure 7A, red color), and in the range between ~27% and ~38% at pH 7.2 (Figure 7A, blue color). On the other hand, coil content is in the range ~42% to ~54% at pH 4.2 (Figure 7B, red color) and between ~33% and 47% at pH 7.2 (Figure 7B blue color). Moreover, we also noticed that in two simulations at pH 4.2 the percentage of α-helix is ~2.0% and ~0.05% respectively, and at pH 7.2 ~3.0 and ~12%, respectively. Overall, these boxplots show that PAP248-286 has more coil and less α-helical content at pH 4.2 compared to pH 7.2.

**Figure 7.**
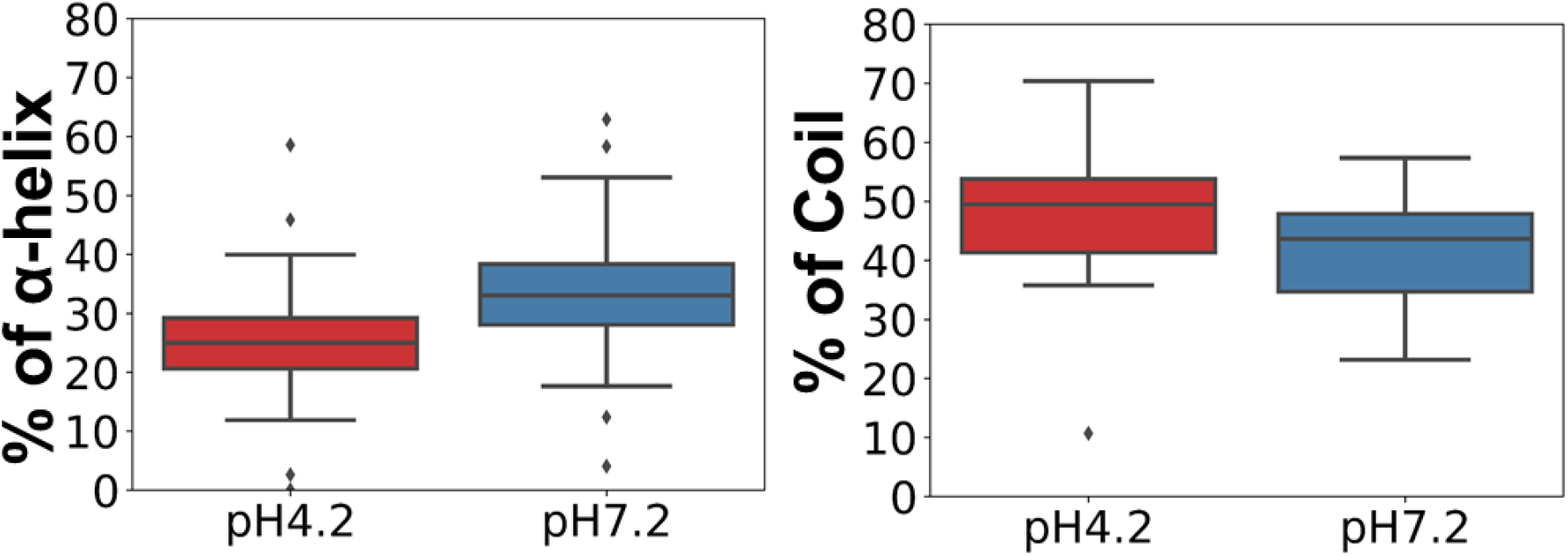
A) Percentage of α-helix content in PAP248-286 at pH 4.2 and pH 7.2, B) Percentage of coil content in PAP248-286 at pH 4.2 and pH 7.2

### Residue contact maps

To identify those residues that make favorable contacts, we computed the residue-residue average contact map and interaction time (see method section for more details). Interestingly, we observed a significant difference in intra-peptide contacts at the two pH values (Figure 8A, B). At pH 4.2, Arg10 and Leu11 in the N-terminal region, Ile20 and Leu21 in the central region, and Met24 in the C-terminal region form contacts with the residues Lys35-Tyr39 for at least 10% of the simulation time (Figure 8C, D). At pH 7.2, only Ile20 and Met24 form contacts with residues Lys35-Tyr39 for 10% or more of the simulation time. Another significant difference in contacts at the two pH values can be observed in the middle region residues Leu16-Ile20 and in the N-terminal residues. At pH 7.2 Leu16 forms contacts with Gln5 and Ser9; Val17 forms contacts with Ile2, Lys6, Ser9, and Arg10; Asn18 interacts with Leu11, while Ile20 forms contacts with residues Ile2, Gln5, Leu11 for 10% or more of the simulation time. However, these contacts are absent at pH 4.2. Moreover, at pH 7.2 the N-terminal residues Gly1 and Gln5 form contacts with each other for ≥ 20% of the simulation time, while at pH 4.2 their interaction time is below 10%. Overall, these data suggest that at pH 7.2 more residues remain within a rcut cut-off range. Additionally, interactions are also formed between the residues for longer periods at pH 7.2 compared to pH 4.2.

**Figure 8.**
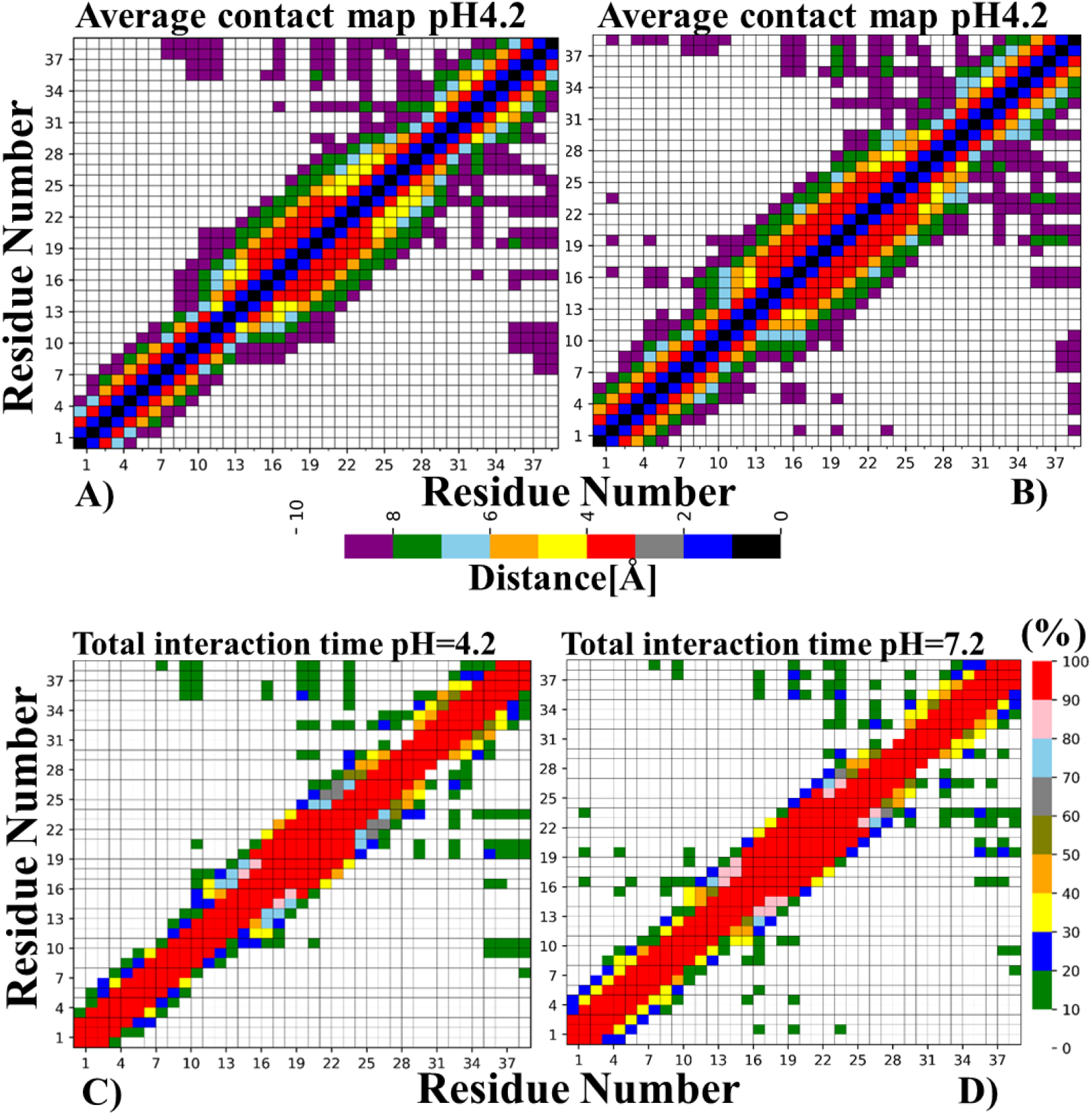
A) Average contact map of PAP248-286 at pH 4.2. B) Average contact map of PAP248-286 at pH 7.2. C) Total interaction time between residues of PAP248-286 at pH 4.2. D) Total interaction time between residues of PAP248-286 at pH 7.2.

### Interaction type

To assess the nature of the interactions that take place between the residues of the PAP248-286 peptide at the two different pH values, we have plotted interaction types between the residues (see methods section for more details) (Figure 9). In the N-terminal region of the peptide there are hydrogen bonding interactions and two salt bridges at both pH values. The first salt bridge is between Lys6 and Glu7, while the second one is between Lys8 and Glu7. In the middle region and in the C-terminal region of PAP248-286, multiple hydrogen bonding and hydrophobic interactions are formed at both pH values. However, we observe no major difference in the interaction pattern between the two pH values, except the presence of a single hydrophobic interaction between Tyr33 and Leu36 at pH 4.2, which is absent at pH 7.2. On the other hand, at pH 7.2 Ala27 and Ile30 form a hydrophobic interaction that is absent at pH 4.2. Overall, hydrogen bonding and hydrophobic interactions are largely prevalent at both pH values (Figure 9).

**Figure 9:**
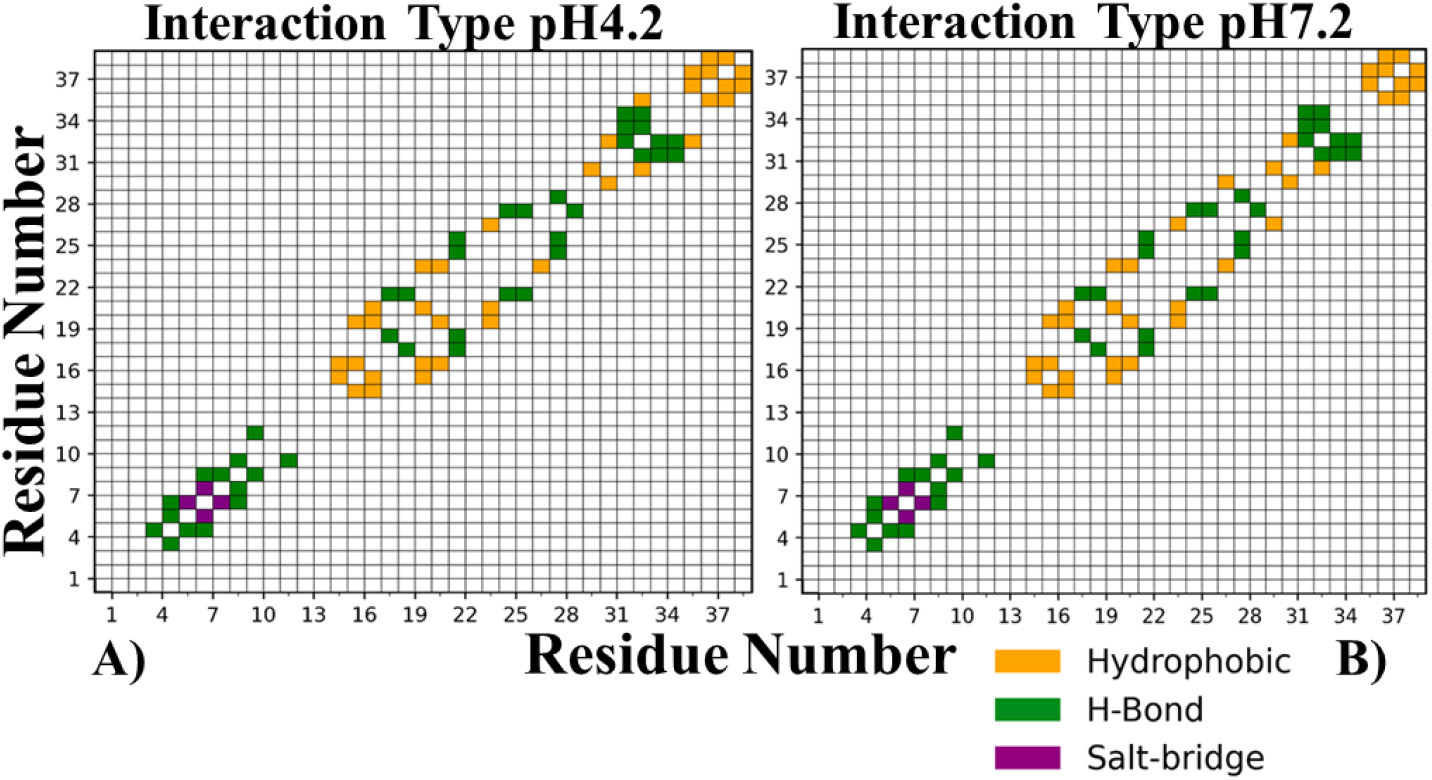
PAP248-286 interaction types A) at pH = 4.2, and B) at pH = 7.2.

### Free energy landscape of PAP248-286

The energy landscape of a protein is the mapping of the different energy levels corresponding to all its possible conformations^50^. To identify the misfolding pathway and the different conformational states adopted by PAP248-286 at pH 4.2 and 7.2, we constructed a twodimensional free energy landscape (FEL) contour map using fraction of native contacts (Q) and RMSD (Å) as reaction coordinates (Figure 10). The free energy landscape of PAP248-286 features three energy basins at pH 4.2 and four energy basins at pH 7.2. At pH 4.2 (Table 1), basin A is in the Q range (~0.869-0.892) and in the RMSD range (~7.16-8.19 Å). In basin A, PAP248-286 contains a long helical region while part of the N- and C-terminal regions of the peptide contain random coil structures. The second basin (B), which is the lowest energy basin (Figure 10A) of PAP248-286, is in the Q range (~0.7389-0.7946) and in the RMSD range (~11.3 to 11.84 Å). In basin B, the N- and C-terminal regions of PAP248-286 are completely unstructured and the central region of the peptide forms a helix. The third energy basin (C) is in the Q range (~0.6079-0.6257) and in the RMSD range (~12.74 to 14.14 Å). In basin C, the structure of PAP248-286 does not contain any helical motif and most of the peptide forms a random coil, while the C-terminal residues Leu36 and Leu37 and the central region residues Gly14 and Val15 form a small β-sheet with an antiparallel arrangement. It is worth noting that the energy barrier between basin B and C is significantly higher than the energy barrier between basin A and B. At pH 7.2, basin A is at Q (0.9151) and RMSD (6.88 Å). The basin A structure features a “U-shaped” topology in a predominantly helical form. Basin B, which is the lowest energy basin at this pH value, is in the Q range (~0.7769-0.7885) and in the RMSD range (~9.86-10.41 Å). Hence, it is highly restricted compared to the lowest energy basin (B) at pH 4.2. In basin B at pH 7.2, the structure of PAP248-286 features an N-terminal random coil region and a long helical portion formed by the central region of the peptide and 6 residues of the C-terminal region. In basin C, which is at Q (0.6756) and RMSD (10.56 Å), the structure of PAP248-286 features N- and C-terminal random coils while part of the central region maintains a helical structure. Basin D is at Q (0.6044) and RMSD (14.12 Å). In this basin, PAP248-286 adopts a random coil and antiparallel β-sheet-rich structure. The antiparallel β-sheet in the structure is formed by the N-terminal residues Arg10, Leu11 and Gln12 and by the C-terminal residues Ser32, Tyr33 and Lys34. The energy barrier between basin C and D is lower than the energy barrier between basin B and C, indicating that the conversion between the C and the D state could be more rapid than the conversion between the B and the C state. Overall, FEL analysis shows that i) there are more local minima at pH 7.2 than at pH 4.2 and ii) basin C at pH 7.2 and the lowest energy structure at pH 4.2 (basin B) are almost identical to the structure of PAP248-286 determined by NMR in membrane environment (PDB id: 2L3H)^17^. iii) PAP248-286 forms antiparallel β-sheet structures at both pH values.

**Figure 10:**
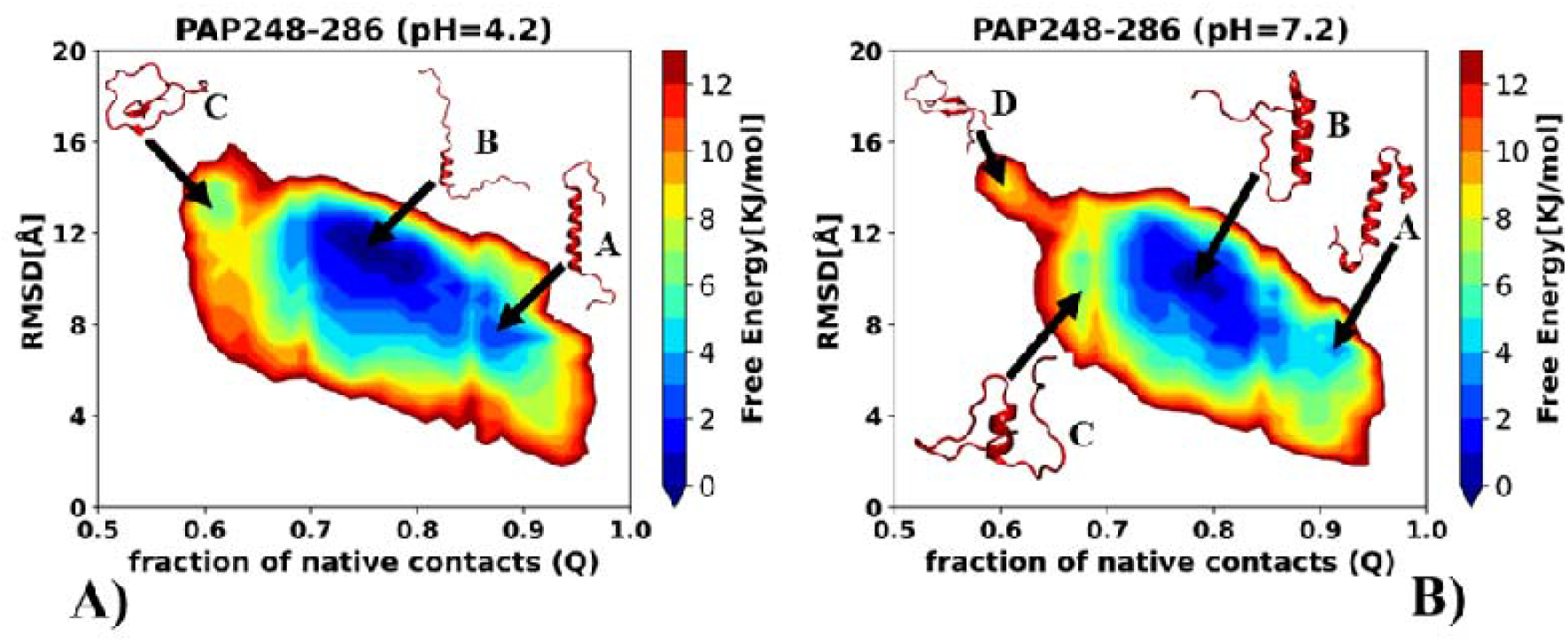
Free energy landscape of PAP248-286 at A) pH=4.2 and B) pH=7.2

**Table 1.**
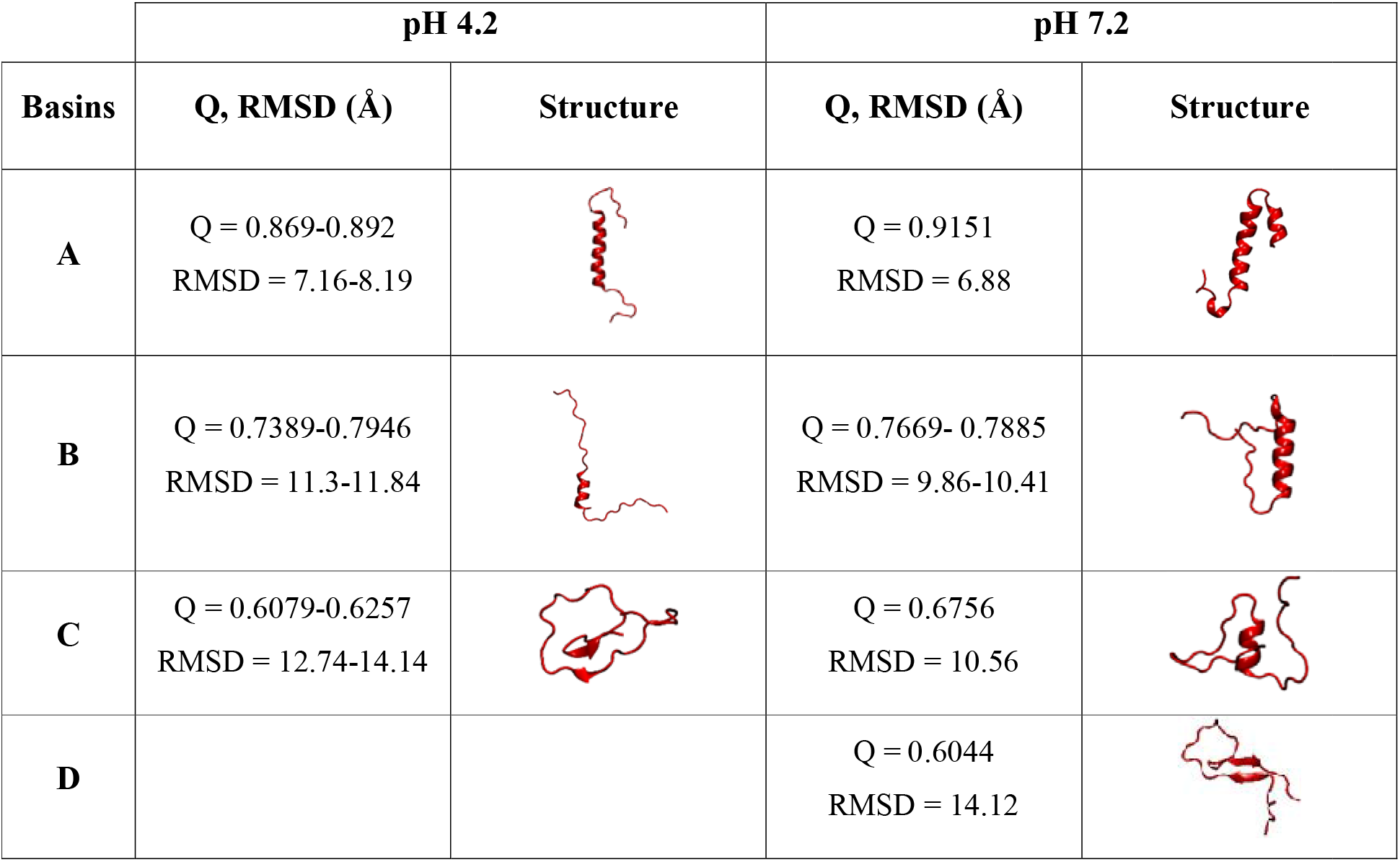
Values of native contacts and RMSD and respective structures for each basin at both pH values.

| Basins | pH 4.2 |  | pH 7.2 |  |
| --- | --- | --- | --- | --- |
|  | Q, RMSD (Å) | Structure | Q, RMSD (Å) | Structure |
| **A** | Q = 0.869-0.892<br>RMSD = 7.16-8.19 |  | Q = 0.9151<br>RMSD = 6.88 |  |
| **B** | Q = 0.7389-0.7946<br>RMSD = 11.3-11.84 |  | Q = 0.7669- 0.7885<br>RMSD = 9.86-10.41 |  |
| **C** | Q = 0.6079-0.6257<br>RMSD = 12.74-14.14 |  | Q = 0.6756<br>RMSD = 10.56 |  |
| **D** |  |  | Q = 0.6044<br>RMSD = 14.12 |  |

### Clustering of Structures

Clustering of protein conformations from multiple simulation trajectories help to divide protein structures into different groups that share similar topological features^51^. To assess the effect of pH, we have performed RMSD-based clustering using 0.6 million structures at both pH 4.2 and pH 7.2 (Figure 11). At both pH values, the top five conformational pools show that PAP248-286 adopts different conformations. At pH 4.2, we observe that cluster-3 (CL3) and cluster-4 (CL4) feature the least α-helical content, while cluster-5 has the most. At pH 7.2, cluster-4 (CL4) and cluster-5 (CL5) show the least α-helical content, while cluster-1 (CL1) displays the most. Interestingly, at both pH values most of the structures adopt a “U-shaped” conformation, which suggests that the PAP248-286 peptide adopts prevalently a “U-shaped” topology. Overall, clustering data suggests that the PAP248-286 peptide adopts significantly different conformations at the two pH values.

**Figure 11.**
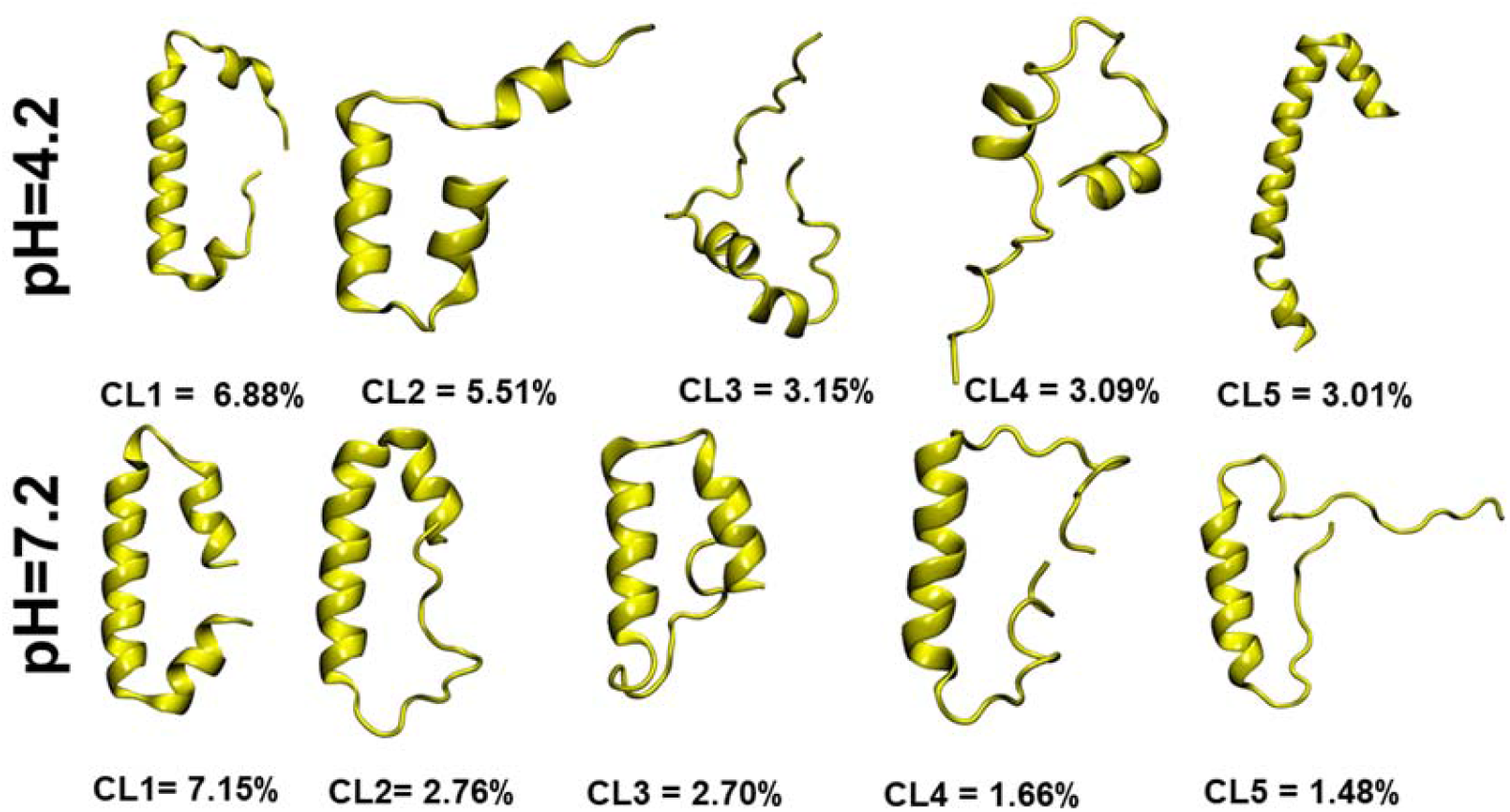
RMSD-based clusters of PAP248-286 at A) pH=4.2 and B) pH=7.2

## Discussion & conclusions

Our MD simulations of the PAP248-286 peptide at two different pH values have shown that more conformational changes take place at pH 4.2 than at pH 7.2. At the acidic pH, less hydrogen bonds and less intra-molecular contacts are formed than at pH 7.2. In addition, we identified a clear difference in the formation of a hydrophobic interaction between a pair of residues at the two pH values, which could be extremely relevant considering that, typically, native and non-native hydrophobic interactions are a crucial driving force for protein secondary structure folding and misfolding processes^52,53^. Furthermore, we also observed significant differences in the free energy landscape of the peptide at the two pH values, featuring more local minima at pH 7.2 than at pH 4.2. Finally, clustering analysis showed that at pH 7.2 the PAP248-286 peptide has more helical content in different clusters than at pH 4.2. Overall, these data suggest that PAP248-286 adopts significantly different conformations at the two physiological pH values. In previous experimental studies, Ramamoorthy & coworkers^7, 14^ showed that the PAP248-286 peptide enhances viral infection *in vivo* in the acidic vaginal environment (pH ~4.2), where it also promotes interaction with lipid bilayers, but not at neutral pH (~7.2). In general, membrane-active peptides are positively charged, amphiphilic in nature and short in length. Moreover, they are commonly unstructured in solution^54^. Based on previous studies and on the data that provided herein, we suggest that the larger extent at which secondary structure content, especially α-helical, and hydrogen bonds are lost at pH 4.2 renders the peptide more flexible. Furthermore, the higher net positive charge of PAP248-286 at pH 4.2 than at pH 7.2 could drive the PAP248-286 peptide to form more interactions with lipid membranes and enhance HIV infection at pH 4.2. We also observed that PAP248-286 adopts a “U”-shaped conformation, which is in line with a previous study by Agrawal *et. al* ^27^ on Aβ peptide, where early stage misfolding of Aβ peptide was shown to feature the formation of a “U-shaped” structure.

In summary, this study provides atomic-level insight into structure dynamics of PAP248-286 at two different pH values and helps unravel the physiological factors that may play an important role in enhancing HIV infection at the acidic vaginal pH value.

## ACKNOWLEDGEMENTS

N.A. acknowledges the ERDF for a postdoctoral grant (No. 1.1.1.2/VIAA/4/20/757). E.P. thanks the ERDF project BioDrug (No. 1.1.1.5/19/A/004) and the Latvian Council of Science (grant No. lzp-2020/2-0013) for financial support. We would like to thank Latvian Institute of Organic Synthesis, Riga for supercomputing resources. N.A. would also like to thank CHPC, Cape Town for supercomputing resources.

